# Palmitoylation of death receptor p75^NTR^ contributes to Alzheimer’s disease progression by regulating APP trafficking and degradation

**DOI:** 10.1101/2025.10.28.685019

**Authors:** Yanchen Ma, Meng Xie, Carlos F. Ibáñez

## Abstract

Although protein palmitoylation has been associated with Alzheimer’s Disease (AD), it remains unclear whether or how palmitoylation of specific proteins contributes to any of the pathological features of AD. The p75 neurotrophin receptor (p75^NTR^) contributes to AD progression by regulating the intracellular trafficking and amyloidogenic processing of amyloid precursor protein (APP). p75^NTR^ is palmitoylated at a juxtamembrane cysteine but it is currently unknown whether this has any effect on its role in AD. Here, we report that 5xFAD mice, an animal model of AD, expressing a palmitoylation-deficient mutant of p75^NTR^ (p75^C281A^) display significantly attenuated neuropathology and cognitive deficits. p75^C281A^ showed enhanced internalization, trafficking to Rab5/Rab7 endosomes and lysosomal-mediated degradation. In mutant p75^C281A^ neurons, APP displayed accelerated co-internalization with p75^NTR^, increased trafficking to late endosomes and lysosome, and enhanced degradation, thereby limiting neuronal Aβ production. Interestingly, the brain of 5xFAD mice shows increased levels of p75^NTR^ palmitoylation. These results indicate that palmitoylation of p75^NTR^ enhances its stability and, indirectly, that of APP by reducing their trafficking to the lysosome, resulting in increased Aβ accumulation and neuropathology in the AD brain. Selective inhibitors of p75^NTR^ palmitoylation may find applications in the treatment of AD

## Introduction

Aberrant aggregation of β-amyloid peptide (Aβ), generated through proteolytic cleavage of amyloid precursor protein (APP) by the sequential actions of β- and γ-secretases, is a primary pathological hallmark of Alzheimer’s Disease (AD) (Karran et al., 2011; Selkoe and Hardy, 2016; Vassar et al., 1999). Familial cases of AD (FAD) are characterized by mutations in the genes encoding APP and presenilin (PSEN), a critical component of γ-secretase, that result in increased generation of Aβ and amyloid plaques, establishing a principal causative link between Aβ production and AD pathogenesis (Selkoe and Hardy, 2016). Interestingly, several of the key proteins associated with AD, including APP, the β-secretase BACE1 and components of the γ-secretase complex, such as nicastrin and APH-1, have been shown to undergo palmitoylation (Cho and Park, 2016), a post-translational lipid modification involving the covalent attachment of palmitate to cysteine residues via a thioester bond. This reversible protein modification plays a crucial role in the dynamic regulation of the stability, trafficking and function of membrane-associated proteins (Peng et al., 2024). Protein palmitoylation is particularly widespread in the nervous system, suggesting important roles in neurobiological processes (Blanc et al., 2015; Holland and Thomas, 2017; Peng et al., 2024). Intriguingly, a recent study reported a marked increase in protein palmitoylation in the hippocampus of 3xTg-AD mice, a common model of AD (Natale et al., 2024). The same study found that treatment of these mice with the broad spectrum palmitoylation inhibitor 2-BP counteracted the progression of AD-related phenotypes. Although these observations suggest a role for protein palmitoylation in AD, it is still unclear whether palmitoylation of specific proteins contributes to any of the pathological features of this disease.

The p75 neurotrophin receptor (p75^NTR^), a member of the “death” receptor superfamily implicated in the regulation of AD pathology, has been shown to be palmitoylated at a cysteine residue in its intracellular juxtamembrane region (Barker et al., 1994). p75^NTR^ functions as a receptor for a family of growth factors known as the neurotrophins, which includes nerve growth factor (NGF) and brain-derived neurotrophic factor (BDNF) (Underwood and Coulson, 2008; Ibáñez and Simi, 2012). Similar to other death receptors, p75^NTR^ can induce cell death as a mechanism for clearing damage produced after a lesion or insult. Following severe injury or disease, however, p75^NTR^ can also amplify tissue damage as a result of overactivation and/or overexpression (Ibáñez and Simi, 2012). In AD, elevated expression of p75^NTR^ has been found in the brains of both patients (Ernfors et al., 1990; Mufson and Kordower, 1992; Hu et al., 2002; Chakravarthy et al., 2012) as well as animal models (Chakravarthy et al., 2010; Wang et al., 2011; Yi et al., 2021). p75^NTR^ can directly interact with APP (Yi et al., 2021; Fombonne et al., 2009) and contributes to AD-associated neuropathology by accelerating Aβ production, neurite degeneration and cognitive impairment in mouse models of AD (Knowles et al., 2009; Yi et al., 2021; Wang et al., 2011). A recent randomized, placebo-controlled phase 2a clinical trial using a small molecule “modulator” of p75^NTR^ found mild measurable improvements in several AD biomarkers as well as cognitive functions, although the latter did not reach statistical significance (Shanks et al., 2024), supporting the use of p75^NTR^ as a therapeutic target in AD.

The physiological significance of p75^NTR^ palmitoylation remains largely unknown. Aside from the initial report (Barker et al., 1994), two later studies using overexpression in cultured cells indicated that palmitoylation of p75^NTR^ may play a role in the ability of the receptor to induce cell death (Underwood et al., 2008) and regulate expression of the *Ncam1* gene (Mirnics et al., 2005). Given the established role of p75^NTR^ in amplifying AD neuropathology, and the association between AD and protein palmitoylation, we set out to investigate whether palmitoylation of p75^NTR^ affects AD progression in the 5xFAD mouse model, one of the most aggressive animal models of AD (Oakley et al., 2006). Unexpectedly, we found that loss of p75^NTR^ palmitoylation was neuroprotective through a mechanism that enhanced APP turnover thereby reducing Aβ production. These results identify p75^NTR^ as one of the specific proteins in which palmitoylation contributes to AD neuropathology.

## Results

### Increased palmitoylation of p75^NTR^ in the hippocampus of 5xFAD mice

In view of a recent report describing increased protein palmitoylation in the hippocampus of 3xTg-AD mice (Natale et al., 2024), we sought to determine whether palmitoylation of p75^NTR^ was altered in 5xFAD mice. Applying the ABE reaction to tissue lysates from 2 month old mouse hippocampus we found elevated levels of p75^NTR^ palmitoylation in hippocampus of 5xFAD mice (Figure 1A). Although p75^NTR^ expression has been shown to be upregulated at later stages of the disease in 5xFAD mice (Yi et al., 2021), at 2 month of age, 5xFAD mice expressed p75^NTR^ at comparable levels to wild type (Figure 1B). These data indicate increased palmitoylation of p75^NTR^ in the hippocampus of 5xFAD mice.

**Figure 1.**
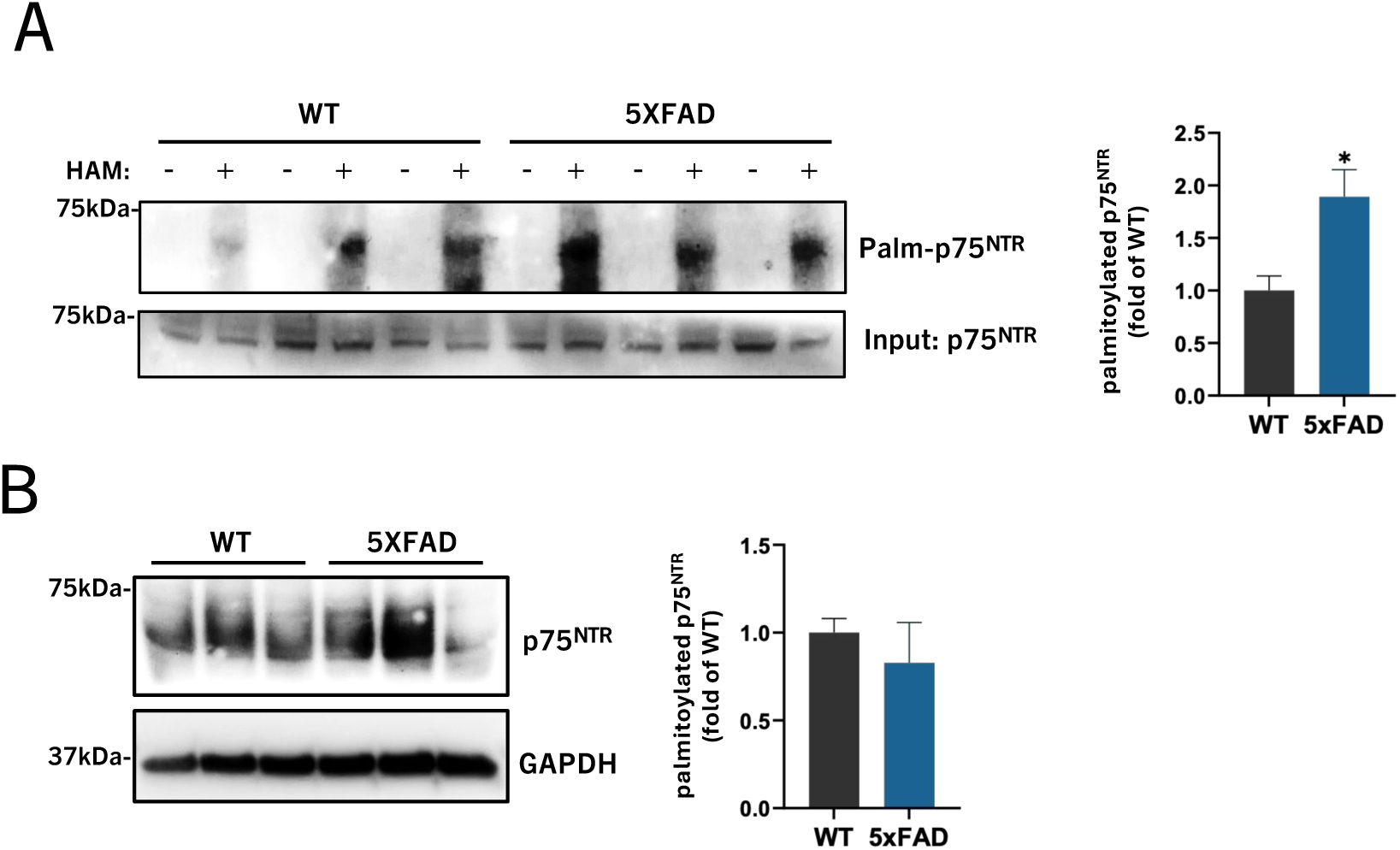
Increased palmitoylation of p75^NTR^ in the hippocampus of 5xFAD mice. (A) ABE assay detection by Western blotting of p75^NTR^ palmitoylation in hippocampus dissected from WT and 5xFAD mice at 2 months of age. Statistical analysis by unpaired t-test; mean ± SEM; N=3 animals per group; *, p<0.05. (B) Western blot of total p75^NTR^ level in the hippocampus of WT and 5xFAD mice at 2 months of age. Statistical analysis by unpaired t-test; mean ± SEM; N=3 animals per group.

### Palmitoylation-deficient p75^NTR^ shows reduced protein half-life and increased susceptibility to lysosomal degradation in hippocampal neurons

In order to investigate the physiological importance of p75^NTR^ palmitoylation for the development of AD, we generated knock-in mutant mice with Ala substituting for Cys at position 281 of the mouse p75^NTR^ protein (herein termed p75^C281A^) (Supplementary Figure S1). Applying the ABE assay to extracts from the hippocampus of these mice, we could verify reduced p75^NTR^ palmitoylation in postnatal day 3 (P3) hippocampus from mutant heterozygous (herein denoted as m/+) and complete ablation in mutant homozygous (denoted m/m) (Figure 2A). In the course of these experiments, we noted that p75^NTR^ was expressed at somewhat lower levels in hippocampal extracts from P7 m/m homozygous mutants compared to wild type mice (Figure 2B) without changes in mRNA expression (Supplementary Figure S2). A parallel decrease was also observed in the levels of p75^C281A^ at the plasma membrane (Figure 2C). We therefore investigated whether the lack of palmitoylation had a destabilizing effect on the p75^NTR^ protein. Indeed, using hippocampal neuron cultures subjected to cycloheximide (CHX) treatment, we observed a reduced protein half-life in mutant p75^C281A^ compared to wild type p75^NTR^ (Figure 2D), suggesting that palmitoylation stabilizes p75^NTR^. In agreement with this, a similar decrease in the half-life of wild type p75^NTR^ protein was observed in hippocampus neuron cultures treated with the palmitoylation inhibitor 2-BP (Figure 2E). We then asked whether C281A mutant p75^NTR^ was being degraded by the proteasome or the lysosome pathways. To this end, hippocampal neuron cultures derived from m/m mouse embryos were treated with the proteasome inhibitor MG132 or the lysosome inhibitor Pepstatin A. We found that the latter, but not MG132, could blunt the degradation of palmitoylation-deficient p75^NTR^ mutant (Figure 2F), indicating that lack of palmitoylation increases the susceptibility of p75^NTR^ to lysosomal degradation.

**Figure 2.**
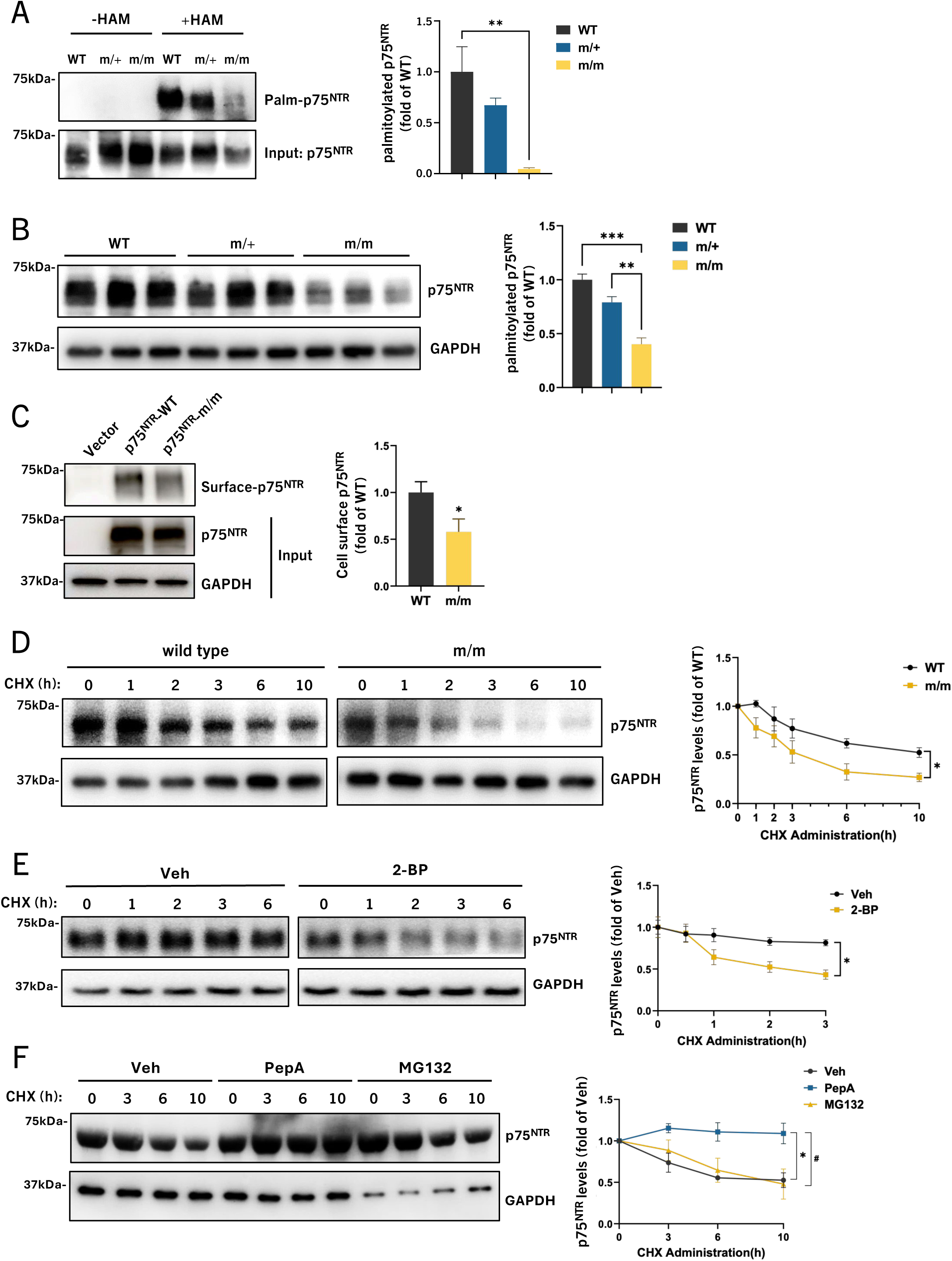
Palmitoylation-deficient p75^NTR^ shows reduced protein half-life and increased susceptibility to lysosomal degradation in hippocampal neurons. (A) ABE assay analysis of p75^NTR^ palmitoylation level in the hippocampus extracts from WT, Cys^281^Ala mutant-heterozygous (m/+) and homozygous (m/m) mice at postnatal day 3. Statistical analysis by one-way ANOVA followed by Tukey’s multiple comparisons test; N=3 animals per group; **, p<0.01. (B) Western blot analysis of protein level of p75^NTR^ in the hippocampus extracts from WT, m/+ and m/m mice at postnatal day7. Statistical analysis by one-way ANOVA followed by Tukey’s multiple comparisons test; mean ± SEM; N=3 animals per group; **, p<0.01; ***, p<0.001. (C) Surface biotinylation assay detection of plasma membrane located p75^NTR^ in HEK293T cells transfected with plasmids expressing either WT or m/m p75^NTR^. Statistical analysis by unpaired t-test; mean ± SEM; N=5 independent experiments; *, p<0.05. (D) Cycloheximide (CHX) analysis of degradation of p75^NTR^ in the cultured hippocampal neurons from WT and m/m mouse embryos. Neurons were cultured 4days in vitro and treated with CHX (50μg/mL) for the indicated times. Statistical analysis by unpaired t-test; mean ± SEM; N=3 independent experiments; *, p<0.05. (E) Western blot analysis of protein level of p75^NTR^ in the cultured hippocampal neurons from WT mouse embryos. Neurons were pretreated with 20 μg/mL 2-Bromopalmitate (2-BP) for 12h before treatment of CHX (50 μg/mL) for the indicated times. Statistical analysis by unpaired t-test; mean ± SEM; N=3 independent experiments; *, p<0.05. (F) Western blot analysis of protein level of p75^NTR^ in the cultured hippocampal neurons dissected from m/m mouse embryos. Neurons were pretreated with 5 μM lysosomal inhibitor Pepstatin A (PepA) or protease inhibitor MG132 for 2h before treatment of CHX (50 μg/mL) for the indicated times. Statistical analysis by one-way ANOVA followed by Tukey’s multiple comparisons test; mean ± SEM; N=3 independent experiments; Veh versus PepA: *p<0.05; Veh versus MG132: #p<0.05.

### Mutant p75^NTR^ lacking palmitoylation shows faster internalization and altered intracellular trafficking in hippocampal neurons

Enhanced lysosomal degradation of the p75^C281A^ protein suggested that the mutant receptor may traffic to lysosomes to a greater extent than wild type p75^NTR^. We therefore investigated possible alterations in the intracellular trafficking of p75^C281A^ in hippocampal neurons. We found that mutant p75^C281A^ internalized from the cell surface of these neurons at a 2-fold higher rate compared to the wild type receptor (Figure 3A). We investigated whether this represented alterations in the association of the mutant receptor with the clathrin or caveolin pathways of internalization and performed co-immunoprecipitation assays in postnatal day 7 (P7) hippocampal extracts from wild type and homozygous m/m mice. We found reduced association of mutant p75^C281A^ with caveolin-1 and a concomitant enhancement of its interaction to β-adaptin, a key component of the clathrin internalization machinery (Figure 3B), suggesting enhanced clathrin-mediated internalization of palmitoylation-deficient p75^NTR^. This is in agreement with the faster internalization of the mutant receptor as it has been proposed that clathrin-mediated internalization of cell surface receptors is faster than caveolin-mediated internalization (Mazumdar et al., 2021). Interestingly, internalized mutant p75^C281A^ showed increased co-localization with Rab5 and Rab7, proteins involved in the trafficking of early and late endosomes, respectively, but lower co-localization with Rab11, which marks endosomes that recycle back to the cell surface (Figure 4A-C), indicating enhanced intracellular trafficking of the mutant receptor. Importantly, mutant p75^C281A^ also showed increased co-localization with Lamp-1, a lysosomal marker (Figure 4D), in agreement with its enhanced susceptibility to lysosomal degradation. Together, these results indicate that palmitoylation of p75^NTR^ enhances the half-life and stability of the protein by increasing the association of the receptor with caveolin at the expense of clathrin, thus slowing its internalization, increasing plasma membrane recycling and reducing p75^NTR^ trafficking to the lysosome.

**Figure 3.**
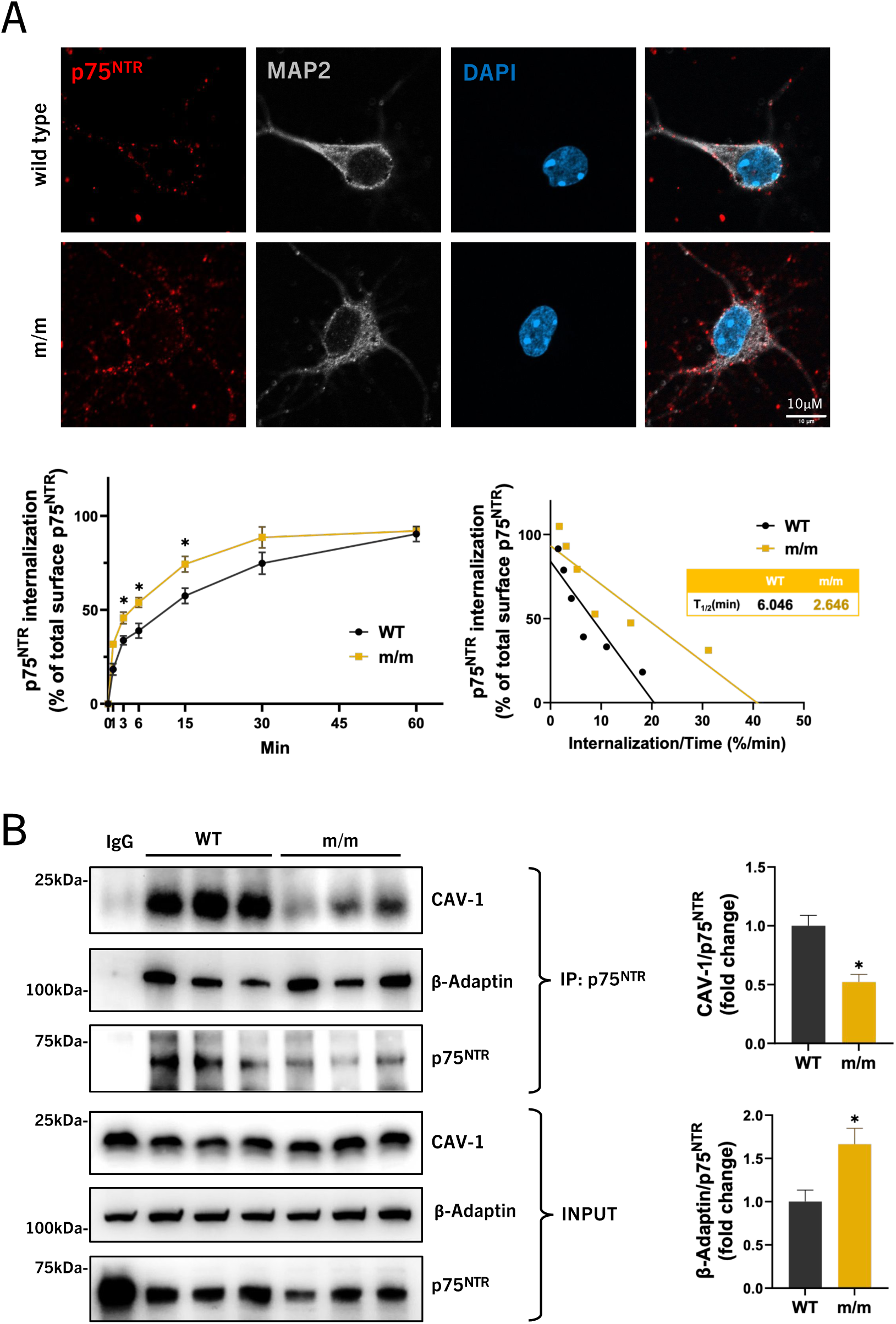
Mutant p75^NTR^ lacking palmitoylation shows faster internalization in hippocampal neurons. (A) Internalization assay of p75^NTR^ in hippocampal neurons from WT and m/m embryos. Microgrpahs show representative images of internalized p75^NTR^ counterstained with microtubule-associated protein 2 (MAP2, dendritic marker for neurons) at 15min of internalization. Graphs below show the kinetics of internalization of p75^NTR^ as in percentage internalization of total surface p75^NTR^ (set to 100%, bottom-left panel) and linear transformation of internalization kinetics (bottom-right panel). T1/2 denotes time for half maximal internalization in minutes. Statistical analysis by two-way ANOVA followed by Šídák’s multiple comparisons test; mean ± SEM; N=3 independent experiments; *, p<0.05. (B) Co-immunoprecipitation of p75^NTR^ with Caveolin-1(CAV-1) and β-Adaptin in the hippocampal extracts from WT or m/m mice at postnatal day7. Statistical analysis by unpaired t-test; mean ± SEM; N=3 animals per group.

**Figure 4.**
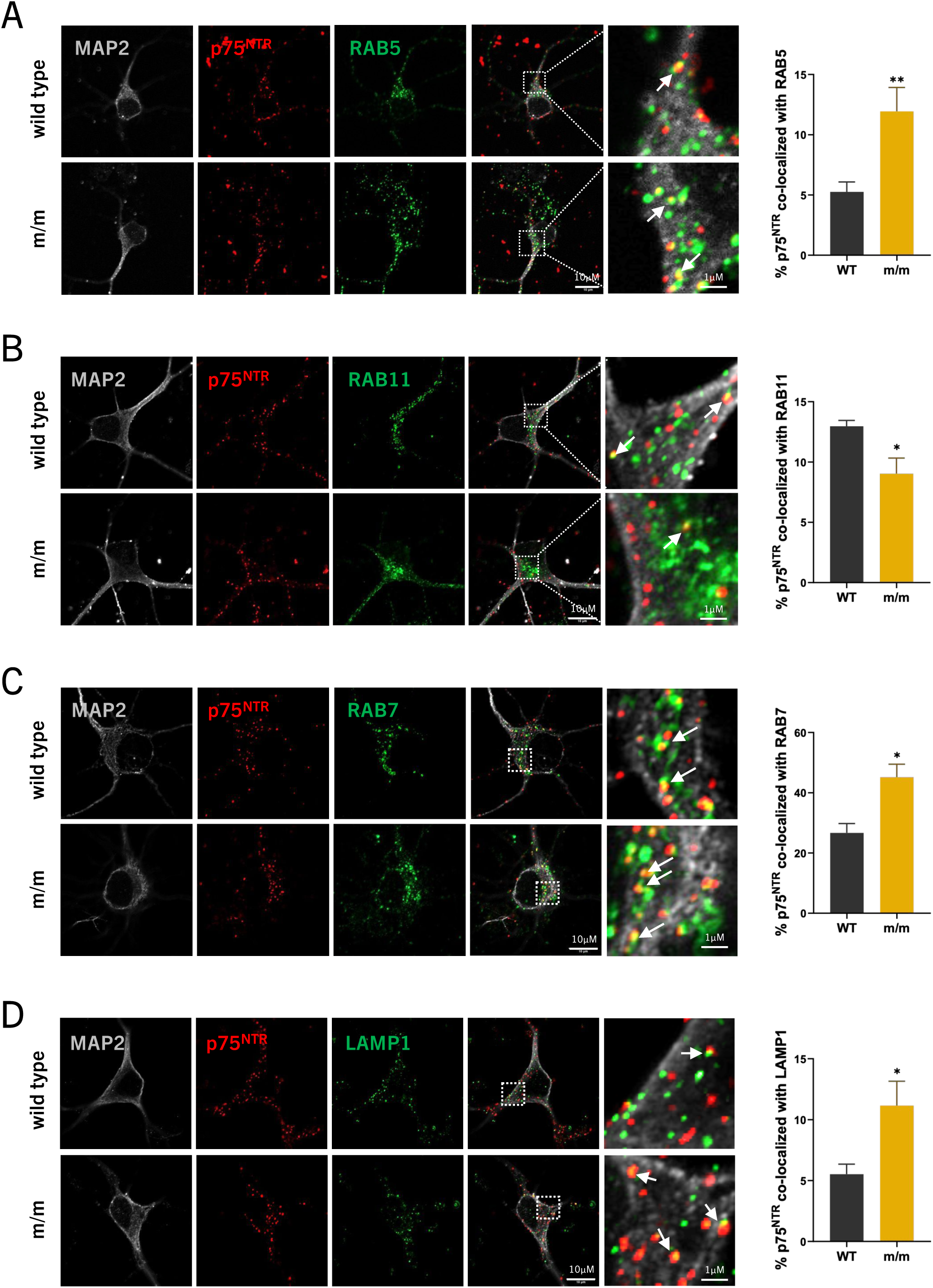
Altered intracellular trafficking of mutant p75^NTR^ lacking palmitoylation. (A) Co-localization of internalized p75^NTR^ with early endosome marker RAB5 in hippocampal neurons after 15 min of internalization. Statistical analysis by unpaired t-test; mean ± SEM; N=3 independent experiments; **p<0.01. (B) Co-localization of internalized p75^NTR^ and recycling endosome marker RAB11 in hippocampal neurons after 30 min of internalization. Statistical analysis by unpaired t-test; mean ± SEM; N=4 independent experiments; *, p<0.05. (C) Co-localization of internalized p75^NTR^ and late endosome marker RAB7 in hippocampal neurons after 60 min of internalization. Statistical analysis by unpaired t-test; mean ± SEM; N=3 independent experiments; *, p<0.05. (D) Co-localization of internalized p75^NTR^ and lysosomal marker LAMP1 in hippocampal neurons after 60 min of internalization. Statistical analysis by unpaired t-test; mean ± SEM; N=4 independent experiments; *, p<0.05.

### Reduced Aβ plaque burden, gliosis and neurite dystrophy in the hippocampus of 5xFAD mice expressing palmitoylation-deficient p75^NTR^

In order to investigate the impact of p75^NTR^ palmitoylation on the progression of AD, p75^C281A^ mutant mice were crossed with 5xFAD mice. As reported previously (Yi et al., 2021), expression of p75^NTR^ became upregulated in the hippocampus of 5xFAD mice compared to wild type controls (Figure 5A). As expected from the reduced stability of palmitoylation-deficient p75^NTR^, the hippocampus of both heterozygous (m/+) and homozygous (m/m) 5xFAD mice expressed lower levels of the mutant receptor, comparable to the levels observed in 5xFAD mice that were heterozygous for a null allele of p75^NTR^ (5xFAD;p75^+/-^) (Figure 5A). In heterozygous form, this null mutation afforded no protection from the accumulation of Aβ plaques in the hippocampus of 5xFAD mice at neither 6 or 9 months (Figure 5B, C). In contrast, Aβ plaque burden was significantly reduced in both m/+ and m/m 5xFAD mice at both ages with a concomitant reduction in plaque size (Figure 5B-D). In agreement with this, the hippocampus of 9 month old mice expressing palmitoylation-deficient p75^NTR^ contained significantly lower levels of Aβ1-42 monomers and oligomers as detected by ELISA (Figure 5E, F). No statistically significant differences were found in the Aβ1-42 levels of hippocampus from 5xFAD;p75^+/-^ mice (Figure 5E, F). This result suggests a protective effect of palmitoylation-deficient p75^NTR^ that can not be accounted for by its reduced expression levels. Intriguingly, Aβ1-42 levels were reduced in hippocampus of heterozygous (m/+) p75^C281A^ 5xFAD mice to a similar extent as in homozygous (m/m) mice (Figure 5E, F). The fact that similar protection was achieved in heterozygous and homozygous p75^C281A^ mutants suggested a dominant effect of palmitoylation-deficient p75^NTR^.

**Figure 5.**
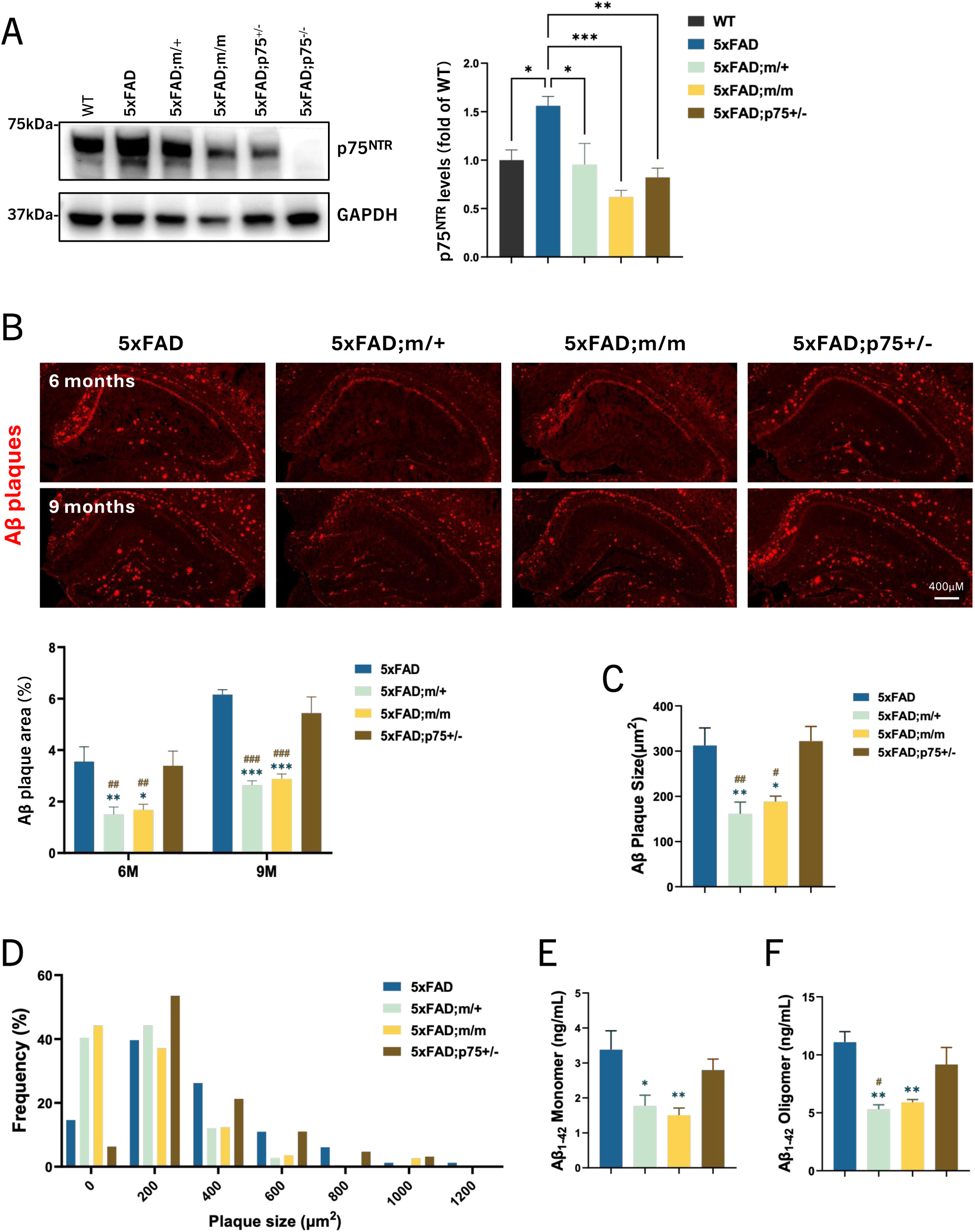
Reduced Aβ plaque burden in hippocampus of 5xFAD mice expressing palmitoylation-deficient p75^NTR^. (A) Western blot analysis of protein level of p75^NTR^ in hippocampal extracts from WT, 5xFAD, 5xFAD homozygous (m/m) and heterzygous (m/+) for palmitoylation-deficient p75^NTR^, and 5xFAD heterozygous knockout for p75^NTR^ (p75^+/-^) mice. Statistical analysis by one-way ANOVA followed by Tukey’s multiple comparisons test; mean ± SEM; N=5 animals per group; *, p<0.05; **, p<0.01; ***, p<0.001. (B) Immunostaining of Aβ plaque (6E10 antibody) in hippocampus of 5xFAD, 5xFAD;m/+, 5xFAD;m/m and 5xFAD;p75^+/-^ mice at 6 months and 9 months of age. Histogram below shows the percentage of hippocampal area occupied by Aβ plaques. Statistical analysis by two-way ANOVA followed by Tukey’s multiple comparisons test; mean ± SEM; N=5 animals per group; *, p<0.05; **, p<0.01; ***, p<0.001 versus 5xFAD; ##, p<0.01; ###, p<0.001 versus 5xFAD;p75+/-. (C) Quantification of average Aβ plaque size in hippocampus of 5xFAD, 5xFAD;m/+, 5xFAD;m/m and 5xFAD;p75^+/-^ mice at 6 months and 9 months of age. Statistical analysis by one-way ANOVA followed by Tukey’s multiple comparisons test; mean ± SEM; N=5 animals per group; *, p<0.05; **, p<0.01 versus 5xFAD; #, p<0.05; ##, p<0.01 versus 5xFAD;p75^+/-^. (D) Frequency distribution analysis for plaque size in hippocampus from 5xFAD, 5xFAD;m/+, 5xFAD;m/m and 5xFAD;p75^+/-^ mice at 9 months of age. (E, F) ELISA determination of Aβ1-42 cotent in hippocampus extracts from 5xFAD, 5xFAD;m/+, 5xFAD;m/m and 5xFAD;p75^+/-^ mice at 9 months of age. Aβ monomers (E) refer to the Tris-buffered saline (TBS) -soluble fraction, while Aβ oligomers (F) refer to the soluble fraction after RIPA buffer extraction of the TBS pellet. Statistical analysis by one-way ANOVA followed by Tukey’s multiple comparisons test; mean ± SEM; N=5 animals per group; *, p<0.05; **, p<0.01 versus 5xFAD; #, p<0.05 versus 5xFAD;p75^+/-^.

In line with reduced Aβ deposition, the hippocampus of m/+;FAD and m/m;FAD presented significantly ameliorated astrogliosis (marked by GFAP expression), microgliosis (marked by Iba1 expression) as well as neurite dystrophy, as revealed by the accumulation of RTN3^+^ labeling around Aβ plaques (Figure 6A-C). The loss of one p75^NTR^ allele in 5xFAD mice had no effect on either gliosis or neurite dystrophy (Figure 6A-C). Again, similar protection was observed in heterozygous and homozygous p75^C281A^ mutants (Figure 6A-C), in line with their reduced Aβ deposition. Together, these data suggest that the specific loss of p75^NTR^ palmitoylation is neuroprotective in AD.

**Figure 6.**
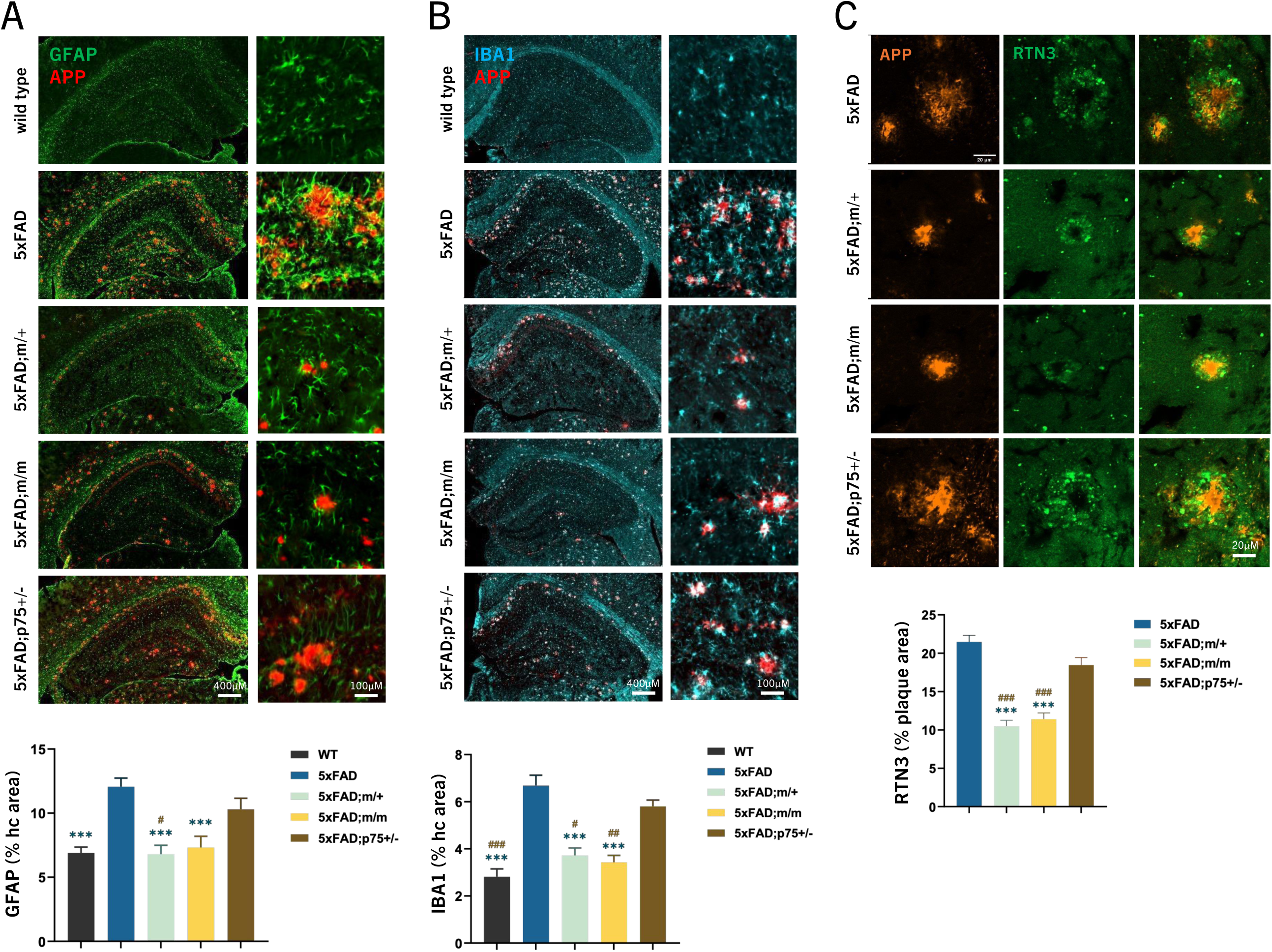
Reduced Aβ gliosis and neurite dystrophy in the hippocampus of 5xFAD mice expressing palmitoylation-deficient p75^NTR^. (A, B) Immunostaining of astrocyte marker glial fibrillary acidic protein (GFAP) (A), microglia marker Ionized calcium binding adaptor molecule1 (Iba1) (B) associated with Aβ plaques in the hippocampus of wild type (WT), 5xFAD, 5xFAD;m/+, 5xFAD;m/m and 5xFAD;p75^+/-^ mice at 9 months of age. Statistical analyses by one-way ANOVA followed by Tukey’s multiple comparisons test are shown below expressed as % of hippocampal (hc) area; mean ± SEM; N=5 animals per group; *, p<0.05; **, p<0.01 versus 5xFAD; #, p<0.05; ##, p<0.01 versus 5xFAD;p75^+/-^. (C) Immunostaining of dystrophic neurite marker reticulon 3 (RTN3) associated with Aβ plaques in hippocampus of 5xFAD, 5xFAD;m/+, 5xFAD;m/m and 5xFAD;p75^+/-^ mice at 9 months of age. Statistical analysis by one-way ANOVA followed by Tukey’s multiple comparisons test is shown below expressed as % of plaque area; mean ± SEM; N=5 animals per group; ***p<0.001 versus 5xFAD; ###, p<0.001 versus 5xFAD;p75^+/-^.

### 5xFAD mice expressing palmitoylation-deficient p75^NTR^ show improved spatial learning and memory

Spatial learning and memory was assessed in 5xFAD mice expressing palmitoylation-deficient p75^NTR^ using the Barnes maze assay. Previous studies have shown that 5xFAD mice present marked deficits in this cognitive test (Yi et al., 2021). As expected, these mice showed little if any learning of the position of the escape tunnel during the training sessions (Figure 7A) and were unable to recognize the target quadrant in the probe session 3 hours after the last training session (Figure 7B). In contrast, both m/+;FAD and m/m;FAD mice learned the position of the escape tunnel at a rate comparable to that of wild type mice (Figure 7A) and recognized the target quadrant above chance during the probe sessions (Figure 7B, C). This result indicates improved cognitive function in the 5xFAD mouse model of AD after the loss of p75^NTR^ palmitoylation.

**Figure 7.**
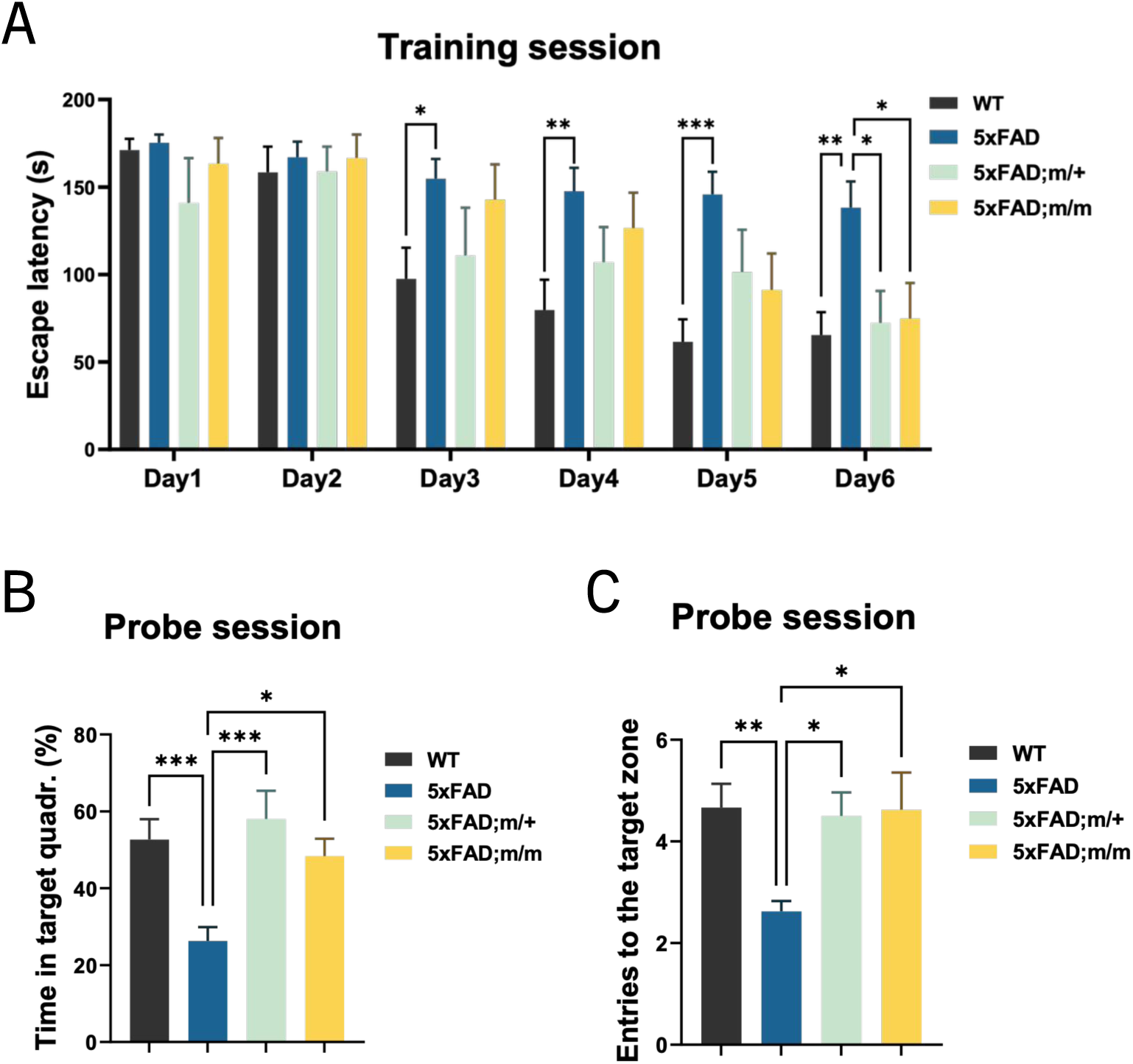
5xFAD mice expressing palmitoylation-deficient p75^NTR^ show improved spatial learning and memory. (A) Escape latency in Barnes maze test of 6-month-old WT, 5xFAD, 5xFAD;m/+ and 5xFAD;m/m mice as indicated. Histograms show mean latency in seconds to find the escape tunnel in 6 consecutive training sessions. Statistical analysis by two-way ANOVA followed by Tukey’s multiple comparisons test; mean ± SEM; N=12 in WT, 16 in 5xFAD, 8 in 5xFAD;m/+ and 5xFAD;m/m; *p < 0.05; **, p<0.01; ***, p<0.001 versus 5xFAD. (B, C) Percentage of time that mice spent in target quadrant (B) and numbers of target hole exploration (C) during the probe trial session 3h after final training of Barnes maze. Statistical analysis by one-way ANOVA followed by Tukey’s multiple comparisons test; mean ± SEM; N=12 in WT, 16 in 5xFAD, 8 in 5xFAD;m/+ and 5xFAD;m/m; *p < 0.05; **, p<0.01; ***, p<0.001 versus 5xFAD.

### Reduced protein half-life, altered intracellular trafficking and decreased amyloidogenic processing of APP in hippocampus of 5xFAD mice expressing palmitoylation-deficient p75^NTR^

Hippocampal extracts from 9 month old 5xFAD mice expressing palmitoylation-deficient p75^NTR^ contained lower levels of full length APP, as well as the CTFβ and sAPPα fragments that result from β- and α-secretase cleavage, respectively, (Figure 8A-C), suggesting enhanced APP protein turn-over in neurons expressing mutant p75^C281A^. In agrement with our histological and behavioral analyses, a similar reduction in APP levels was observed in homozygous (m/m) and heterozygous (m/+) mutants (Figure 8A-C), in line with a dominant effect of palmitoylation-deficient p75^NTR^. Given that APP and p75^NTR^ can directly interact and co-internalize in hippocampal neurons (Yi et al., 2021), we investigated the possibility that palmitoylation-deficient p75^NTR^ affected the stability, trafficking and processing of APP. Human 3xFAD APP (carrying the same three mutations present in 5xFAD mice) expressed in cultured hippocampal neurons by viral transduction showed a faster decay upon CHX treatment in homozygous (m/m) p75^C281A^ neurons compared to wild type cells (Figure 8D), indicating a reduced half-life of 3xFAD APP in neurons expressing palmitoylation-deficient p75^NTR^. The reduced half-life of 3xFAD hAPP in mutant neurons could be ameliorated by treatment with pepstatin A (Figure 8E), indicating lysosomal involvement. In agreement with this, 3xFAD hAPP showed stronger co-localization with the lysosomal marker Lamp-1 in m/m neurons compared to wild type (Figure 8F). A decrease in 3xFAD APP half life was also observed in heterozygous (m/+) neurons (Figure 8D), indicating a dominant effect of palmitoylation-deficient p75^NTR^ on the stability of APP.

**Figure 8.**
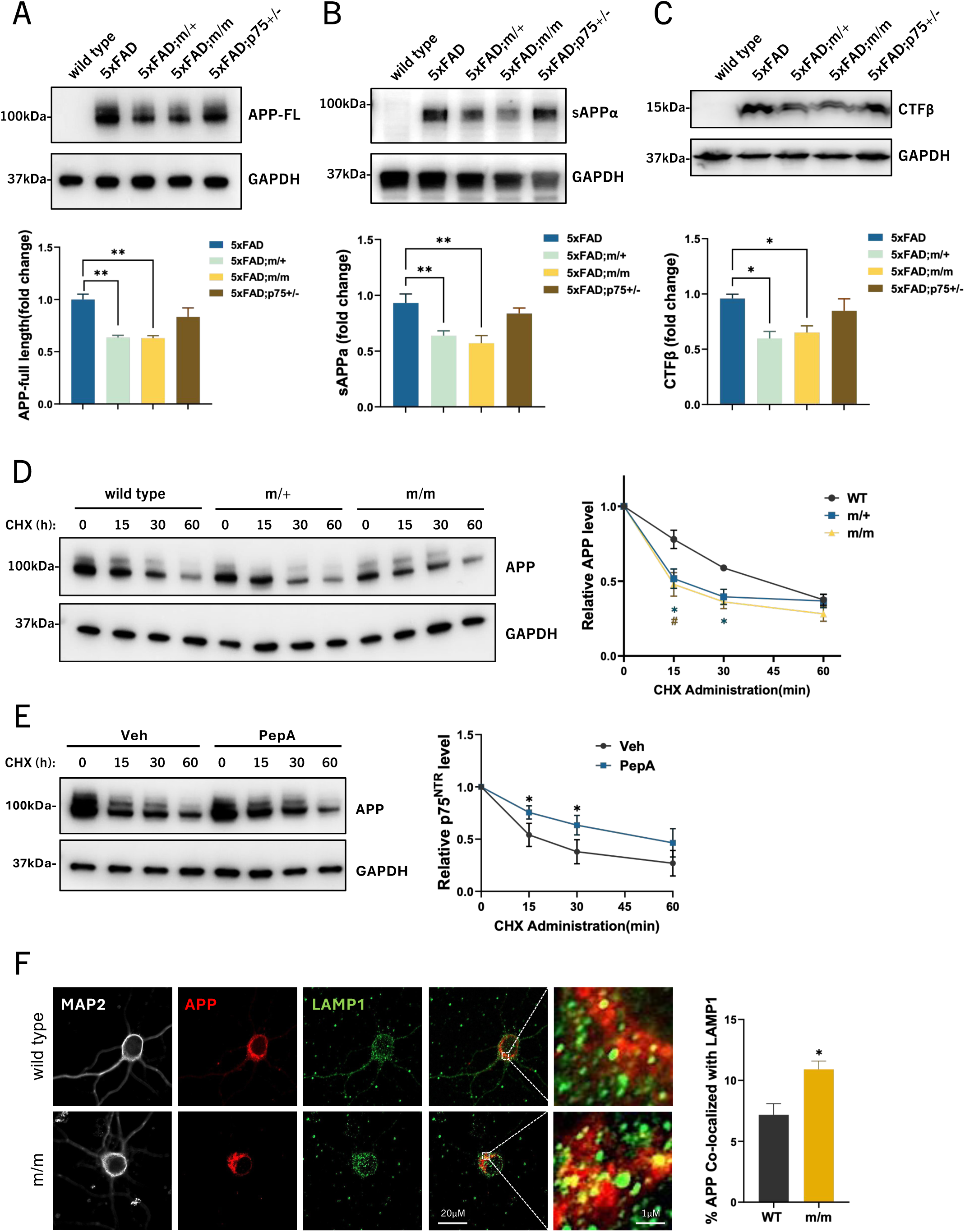
Reduced protein half-life and increased lysosomal localization of APP in AD hippocampal neurons expressing palmitoylation-deficient p75^NTR^. (A) Western blot analysis of full-length APP (6E10 antibody) in the RIPA-soluble fraction of hippocampal extracts from 9-month-old WT, 5xFAD, 5xFAD;m/+ and 5xFAD;m/m mice. Statistical analysis by one-way ANOVA followed by Tukey’s multiple comparisons test is shwon below; mean ± SEM; N=5 in each group. (B) Western blot analysis of soluble APP-α (sAPPα, 6E10 antibody) in the TBS-soluble fraction of hippocampus extract from 9-month-old WT, 5xFAD, 5xFAD;m/+ and 5xFAD;m/m mice. Statistical analysis by one-way ANOVA followed by Tukey’s multiple comparisons test is shown below; mean ± SEM; N=5 in each group. (C) Western blot analysis of C-terminal fragment-β (CTFβ, 6E10 antibody) in the RIPA-soluble fraction of hippocampus extract from 9-month-old WT, 5xFAD, 5xFAD;m/+ and 5xFAD;m/m mice. Statistical analysis by one-way ANOVA followed by Tukey’s multiple comparisons test is shwon below; mean ± SEM; N=5 in each group. (D) Cycloheximide (CHX) pulse-chase assay of APP half-life in wild type (WT), heterozygous (m/+) and homozygous (m/m) p75^NTR^ mutant hippocampal neurons expressing triple mutant human APP introduced by AAV infection. AAV-DJ-3xFAD-APP virus was added to cultured hippocampal neurons dissect from WT, m/+, m/m E17.5 embryos at DIV2 (days in vitro). At DIV7, neurons were treated with CHX (50μg/ml) for indicated time. Statistical analysis by two-way ANOVA followed by Tukey’s multiple comparisons test (right); mean ± SEM; N=4 independent experiments; *, p < 0.05 m/m versus WT; #, p<0.05 m/+ versus WT. (E) Cycloheximide (CHX) pulse-chase assay of APP half-life in homozygous (m/m) p75^NTR^ mutant hippocampal neurons expressing triple mutant hAPP introduced by AAV-DJ-3xFAD-APP virus infection. Neurons were pretreated with lysosomal inhibitor Pepstatin A (5μΜ, 2h) before CHX administration. Statistical analysis by two-way ANOVA followed by Tukey’s multiple comparisons test (right); mean ± SEM; N=3 independent experiments; *p < 0.05. (F) Co-immunostaining of APP with lysosome marker Lysosomal-Associated Membrane Protein 1 (LAMP1) in AAV-DJ-3xFAD-APP infected wild type (WT) and homozygous (m/m) p75^NTR^ mutant hippocampal neurons. Histogram (right) shows the percentage of area of APP that co-localized with LAMP1. Scale bar, 20μm. Statistical analysis by unpaired t-test; mean ± SEM; N=3 independent experiments, ≥15 cells per group in each experiment; *p < 0.05.

Intrigued by the dominant effects of the C281A mutation, we investigated its molecular mechanism by co-transfecting wild type and C281A mutant p75^NTR^ in HEK293 cells followed by assessment of their interaction, palmitoylation and stability. By co-immunoprecipitation, we found that wild type and palmitoylation-deficient mutant p75^NTR^ were able to form homo- and hetero-dimers to a similar extent (Supplementary Figure S3A). Interestingly, palmitoylation of wild type p75^NTR^ was not significantly affected upon co-expression of palmitoylation-deficient mutant (Supplementary Figure S3B), suggesting independent palmitoylation of the two protomers in the p75^NTR^ dimer. Importantly, however, we found that the half-life of wild type p75^NTR^ was significantly reduced upon co-expression of palmitoylation-deficient mutant (Supplementary Figure S3C) indicating a dominant effect of the latter upon the stability of the wild type protomer in the receptor dimer. This result established a plausible mechanism by which the stability of APP may have been affected in m/+ heterozygous neurons.

Finally, we assessed 3xFAD hAPP co-internalization with p75^NTR^ and its intracellular trafficking. Using the Proximity Ligation Assay (PLA), we first verified that wild type and C281A mutant p75^NTR^ interacted with 3xFAD APP to a similar extent (Supplementary Figure S4). In agreement with the enhanced internalization of palmitoylation-deficient p75^NTR^, 3xFAD APP co-internalized with mutant p75^C281A^ to a greater extent than with wild type p75^NTR^ in hippocampal neurons (Figure 9A, B). Moreover, 3xFAD APP/p75^C281A^ co-internalized complexes were present in association with late endosomes marked by Rab7 at significantly higher levels than APP co-internalized with wild type p75^NTR^ (Figure 9B). Importantly, in mutant p75^C281A^ hippocampal neurons, human 3xFAD APP interacted with BACE1 to a lower extent than in wild type neurons as detected by PLA (Figure 9C). In agreement with this, mutant p75^C281A^ neurons expressing human 3xFAD APP released lower levels of Aβ_1-42_ amyloid peptide to the culture medium than neurons expressing wild type p75^NTR^ (Figure 9D). Together, these results suggest that the stabilizing effect of palmitoylation on the p75^NTR^ receptor leads to enhanced APP stability, increased APP interaction with BACE1 and elevated levels of Aβ_1-42_ amyloid peptide, thereby contributing to the development of AD neuropathology.

**Figure 9.**
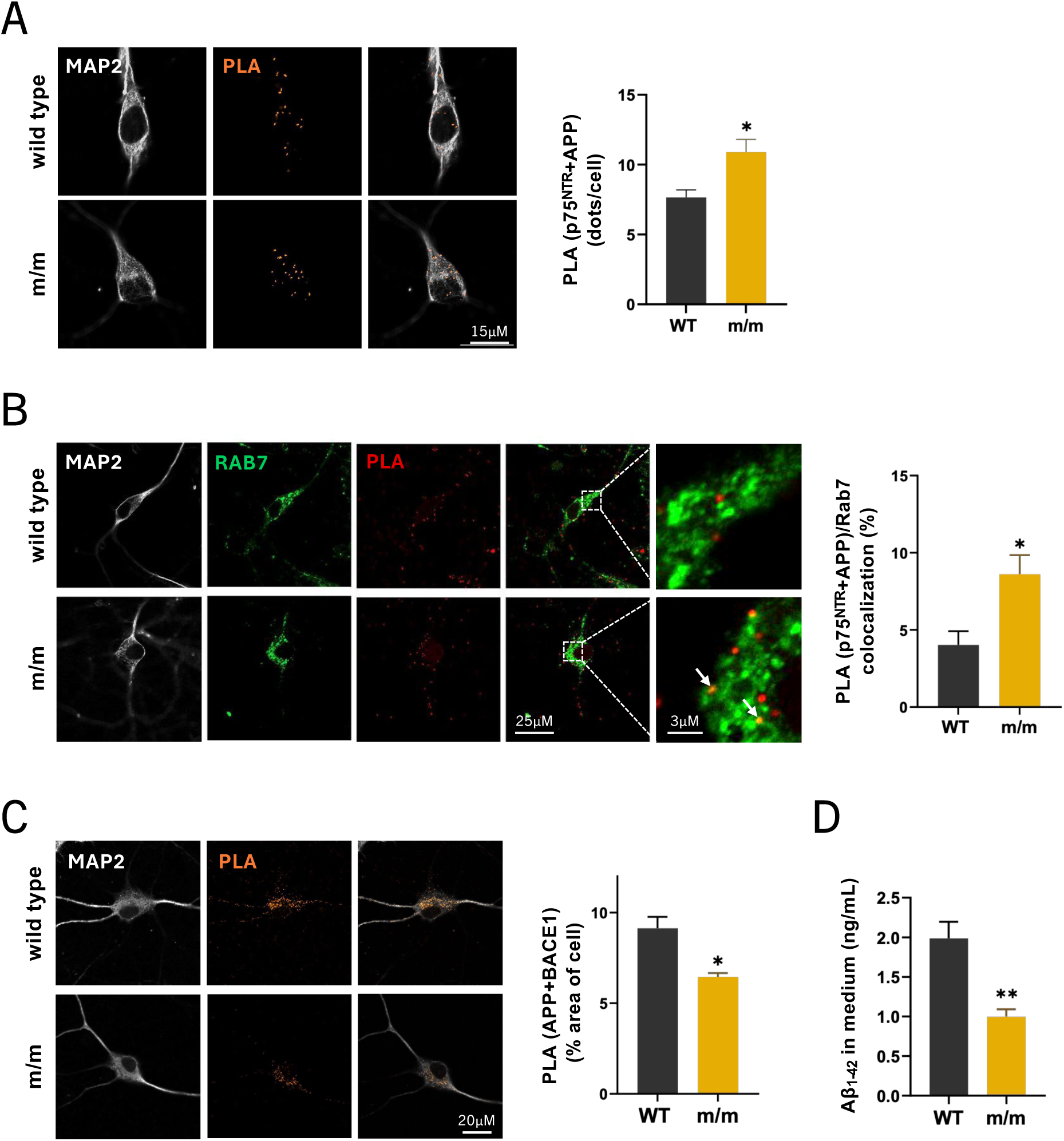
Faster internalization, altered intracellular trafficking and increased amyloidogenic processing of APP in AD hippocampal neurons expressing palmitoylation-deficient p75^NTR^. (A) Proximity-ligation assay (PLA) between 3xhAPP and p75^NTR^ in wild type (WT) and homozygous (m/m) p75^NTR^ mutant hippocampal neurons infected with AAV-DJ-3xFAD-APP virus following 60 min internalization. Histogram (right) shows the numbers of PLA signals per cell. Statistical analysis by unpaired t-test; mean ± SEM; N=3 independent experiments, ≥15 cells per group in each experiment; *p < 0.05. (B) Co-localization of APP/p75^NTR^ co-internalized complexes (detected by PLA) with late endosome/lysosome marker RAS-associated binding protein 7 (RAB7) in wild type (WT) and homozygous (m/m) p75^NTR^ mutant hippocampal neurons infected with AAV-DJ-3xFAD-APP virus. Histogram shows the percentage of area of PLA signals that co-localized with RAB7. Statistical analysis by unpaired t-test; mean ± SEM; N=3 independent experiments, ≥15 cells per group in each experiment; *p < 0.05. (C) Proximity-ligation assay (PLA) between 3xhAPP and BACE1 in wild type (WT) and homozygous (m/m) p75^NTR^ mutant hippocampal neurons infected with AAV-DJ-3xFAD-APP virus. Histogram shows the percentage of area of PLA signals. Statistical analysis by unpaired t-test; mean ± SEM; N=3 independent experiments, ≥15 cells per group in each experiment; *p < 0.05. (D) ELISA detection of Aβ_1-42_ secretion in the conditional medium of cultured hippocampal neurons from wild type (WT) or homozygous (m/m) p75^NTR^ mutant hippocampal neurons infected infected with AAV-DJ-3xFAD-APP virus for 4 days. Statistical analysis by unpaired t-test; mean ± SEM; N=5 independent experiments performed in triplicate; *p < 0.05.

## Discussion

The results of the present study indicate that one key functional consequence of the palmitoylation of p75^NTR^ is a marked increase in its protein stability, achieved primarily by reduced receptor internalization and intracellular trafficking to lysosomes. One consequence of this is that other transmembrane proteins with an ability to interact and co-internalize with p75^NTR^, such as APP, become similarly affected. Based on the hydrophobic of the palmitate tail, palmitoylation has been considered as a mechanism for driving proteins into cholesterol-rich membrane micro-domains known as lipid rafts (Levental et al., 2010; Diaz-Rohrer et al., 2014). Previous reports found small amounts (≈1%) of p75^NTR^ in light-density detergent-resistant fractions, an operational definition of lipid rafts, of transfected cell lines (Huang et al., 1999; Underwood et al., 2008) and rat brain tissue lysates (Gil et al., 2007). Using similar methods, we could only detect minute amounts (<0.1%) of p75^NTR^ in lipid raft fractions of cultured cortical neurons and extracts of 9 month old 5xFAD hippocampus (Supplementary Figure S5A, B). hAPP was also present at very low levels in hippocampal lipid rafts fractions (<1%) (Supplementary Figure S5B). Importantly, we could not detect any significant difference in the levels of hAPP in lipid raft fractions of palmitoylation-deficient p75^NTR^ (m/m) mutant compared to wild type hippocampus (Supplementary Figure S5C, D). These results suggests that in brain neurons expressing physiological levels of p75^NTR^, the receptor may not be present at significant amounts in lipid rafts. The fact that lipid raft levels of 5xFAD hAPP were comparable in m/m and wild type hippocampus further suggests that palmitoylation-deficient p75^NTR^ does not affect hAPP metabolism by altering its localization in lipid rafts. Intriguingly, wild type p75^NTR^ showed higher interaction than the palmitoylated-deficient receptor for caveolin-1, a well-known lipid raft marker. However, caveolins can also be found outside rafts and in non-caveolar regions, as well as in cells that lack caveolae altogether, where they have scaffolding functions that contribute to protein and lipid distribution as well as regulation of the actin cytoskeleton (Head and Insel, 2007; Rezola et al., 2021).

It has been reported in sympathetic neurons that internalized wild type p75^NTR^ evades the lysosomal route at the level of early Rab5^+^ endosomes, instead accumulating in Rab11^+^ vesicles and multivesicular bodies (Escudero et al., 2014). Our present findings, showing preferential localization of internalized palmitoylation-deficient p75^NTR^ in Rab7^+^ endosomes and the lysosome, indicate that the palmitate tail is a key determinant of p75^NTR^ intracellular trafficking. It is possible that palmitate contributes to the partition of internalized p75^NTR^ to distinct endocytic vesicles based on its differential affinity for specific membrane lipid components. It has also been reported that caveolin-containing vesicles are able to avoid lysosomal degradation (Liu et al., 2020), so the lower interaction of palmitoylation-deficient p75^NTR^ with caveolin-1 may in part explain its enhanced lysosomal localization and degradation compared to the wild type receptor.

Previous studies have shown that p75^NTR^ may internalize through both clathrin-dependent and clathrin-independent pathways to different extents depending on the cell-type and the degree of receptor activation (Bronfman et al., 2003; Escudero et al., 2014; Hibbert et al., 2006; Deinhardt et al., 2007). Until recently, APP endocytosis was thought to proceed solely through the clathrin-dependent pathway. However, a recent study reported that APP internalizes through a clathrin-independent mechanism in the somato-dendritic compartment of mouse hippocampal and cortical neurons as well as induced human neurons (Aow et al., 2023). Our observation of a stronger interaction of palmitoylation-deficient p75^NTR^ with β-adaptin, a key component of the clathrin machinery, suggests that, through its association with APP, the mutant receptor may preferentially alter APP endocytosis from somato-dendritic compartments potentially affecting Aβ generation.

In a previous study, we reported that signaling-deficient mutants of p75^NTR^ (i.e. ΔDD and C259A) show reduced receptor internalization, resulting in reduced APP internalization, Aβ generation and neuropathology in the 5xFAD mouse model (Yi et al., 2021). Thus, the observation that palmitoylation-deficient p75^NTR^ shows enhanced internalization and yet is neuroprotective in 5xFAD mice seemed at first paradoxical. However, a key difference resides in the respective fates of the internalized mutant receptors. While internalized palmitoylation-deficient p75^NTR^ readily transits to Rab5^+^ and Rab7^+^ endosomes and subsequently lysosomes, internalized signaling-deficient p75^NTR^ mutants show enrichment in recycling Rab11^+^ endosomes at the expense of Rab7 endosomes (Yi et al., 2021). Due to the low abundance of BACE1 in recycling endosomes, as well as their unfavorable pH conditions for BACE1 activity, lower levels of APP were co-localized with BACE1 and less Aβ was generated in neurons expressing signaling-deficient p75^NTR^ mutants (Yi et al., 2021). Thus, despite having opposite internalization behaviors, signaling-deficient and palmitoylation-deficient p75^NTR^ mutants lead to similar outcomes in the context of AD.

In conclusion, our results indicate that palmitoylation of p75^NTR^ enhances its stability and plasma membrane levels by slowing its internalization and subsequent intracellular trafficking to the lysosome. In doing so, native, palmitoylated p75^NTR^ also stabilizes APP, thereby indirectly contributing to Aβ production and AD pathology. The fact that m/+ heterozygous mutants showed beneficial effects comparable to those observed in m/m homozygous suggests that even moderate inhibition of p75^NTR^ palmitoylation may be sufficient to ameliorate cognitive decline in AD.

## Methods

### Mice

Mice were housed in a 12h light-dark cycle and fed a standard chow diet. The mouse lines utilized in this study have been described previously and are as follows: 5xFAD (Oakley et al., 2006); p75^NTR^ exon 3 knock-out (Lee et al., 1992). The C281A p75^NTR^ knock-in mouse line was generated using CRISPR-Cas9 system by Genetic Manipulation Core in Chinese Institute for Brain Research. All animal experiments were performed under the guidelines set by the Institutional Animal Care and Use Committee of Chinese Institute for Brain Research, Beijing (CIBR-IACUC-028).

### Primary culture of hippocampal neurons

Pregnant female mice were euthanized at 17.5 day of gestation by isoflurane. Hippocampi were aseptically dissected and digested in digestion buffer (10 units/mL Papain, 5 mM L-cysteine and 1000 units DNase I in L-15 medium) for 30 min at 37℃. The digested hippocampal tissue was neutralized with complete medium containing 10% heat-inactivated horse serum then triturated 15 times using a 1 mL pipette tip to dissociate it into a single-cell suspension. Neurons were transferred to coverslips or well plates coated with 0.01% poly-D-lysine (Sigma-Aldrich) and 1 μg/ml laminin (Sigma-Aldrich) followed by one complete medium change after 1hr of seeding. Cultured neurons were maintained in serum-free Neurobasal medium supplemented with B27 (Invitrogen), GlutaMAX (Invitrogen) at 37℃ in 5% CO2.

### Acyl-Biotin Exchange (ABE) Assay

ABE assays were performed according to a previous report (Ma et al., 2022). Briefly, transfected HEK293T cells or hippocampi from postnatal day 3 pups were lysed in 50 mM HEPES pH 7.2, 2% (w/v) SDS, 1 mM EDTA plus protease inhibitors (MCE). 20 Mm methyl-methane thiosulfonate (MMTs) were added to block free thiols at 42℃ for 1hr. Unreacted MMTs were removed by acetone precipitation and pellets were resuspended in buffer containing 0 mM HEPES pH 7.2, 4% (w/v) SDS, 1 mM EDTA. Samples were diluted in buffer containing 0.7 M hydroxylamine (+HAM) or 50mM Tris-HCl (pH7.4, -HAM) with sulfhydryl-reactive HPDP-biotin and incubated at room temperature for 1hr with gentle agitation. Samples were acetone precipitated again to remove HAM and HPDP-Biotin and pellets were resuspended in 4% SDS. Samples were diluted to 0.1% SDS and biotinylated proteins in samples were affinity-purified using neutravidin-conjugated beads at room temperature for 1h. 50mM DTT was used to cleave HPDP-biotin and release purified proteins. The released proteins were denatured in SDS sample buffer and processed for SDS-PAGE.

### Co-Immunoprecipitation, Surface Biotinylation and Western blotting

Hippocampi dissected from postnatal day 7 mice or cell samples were lysed in Co-IP lysis buffer (50mM Tris-HCl pH 7.2, 0.15 NaCl, 0.5mM EDTA, 250 mM sucrose, 60 mM Octyl-β-glucoside, 1% Triton X-100) at 4℃ for 30 min. Cell extract was incubated with p75NTR antibody (Abcam, 8 ug/ml) or HA antibody (CST, 3ug/ml) overnight at 4 ℃. 25ul protein A/G magnetic beads (MCE) were added to lysate and incubated at room temperature for 1hr. After incubation, the beads were washed by PBST buffer for 4 times. The binding proteins were released and denatured in DTT-containing SDS sample buffer and processed for SDS-PAGE.

HEK293T cells plated on 10 cm plate were transfected with HA-tagged p75NTR constructs or control vector. After 24 hr of transfection, the medium was removed and cells were washed twice by 10 ml ice-cold PBS buffer. 6 ml cold cell-impermeant biotin buffer (0.5mg/ml Sulfo-NHS-SS-Biotin, 10 mM MgCl_2_, 0.5% BSA in DMEM without serum or antibiotics) was added to each plates, and incubated at 4 ℃ with gentle agitation. Following incubation, the labeling solution was removed, and cells were incubated in 8.5 ml quenching solution (PBS pH8.0, 10 mM Tris, 100 mM glycine) at 4 ℃ for 10 min. Cells were washed by 5ml PBS for twice and lysed in 500ul lysis buffer (the same as in Co-IP assay) containing protease inhibitors. Lysates were then incubated with 25ul neutravidin-conjugated beads at RT for 2h. After incubation, the biotin-labeled surface protein were released and denatured in DTT-containing SDS sample buffer for further western blot analysis.

Tissue or cell lysates were denatured in DTT-containing SDS sample buffer (Solarbio) and boiled at 95℃ for 5 min. Samples were shortly centrifugated before electrophoresis on polyacrylamide gels. Proteins were blotted on polyvinylidene fluoride (PVDF) membranes (0.22 μm, Millipore). Membranes were blocked by 5% BSA in TBST for 1 hr at room temperature and incubated with primary antibodies overnight at 4 ℃. After washing by TBST for 3 times, membranes were incubated with horseradish peroxidase (HRP) conjugated secondary antibodies (1:10,000, CST) for 1 hr at RT. Immunoblots were then developed using ECL Reagent (Millipore) and exposed by Amersham ImageQuant™ 800 (Cytiva). ImageJ was used to analyze the images.

### Surface Biotinylation

HEK293T cells plated on 10 cm plate were transfected with HA-tagged p75NTR constructs or control vector. After 24 hr of transfection, the medium was removed and cells were washed twice by 10 ml ice-cold PBS buffer. 6 ml cold cell-impermeant biotin buffer (0.5mg/ml Sulfo-NHS-SS-Biotin, 10 mM MgCl2, 0.5% BSA in DMEM without serum or antibiotics) was added to each plates, and incubated at 4 ℃ with gentle agitation. Following incubation, the labeling solution was removed, and cells were incubated in 8.5 ml quenching solution (PBS pH8.0, 10 mM Tris, 100 mM glycine) at 4 ℃ for 10 min. Cells were washed by 5ml PBS for twice and lysed in 500ul lysis buffer (the same as in Co-IP assay) containing protease inhibitors. Lysates were then incubated with 25ul neutravidin-conjugated beads at RT for 2h. After incubation, the biotin-labeled surface protein were released and denatured in DTT-containing SDS sample buffer for further western blot analysis.

### Western blotting

Tissue or cell lysates were denatured in DTT-containing SDS sample buffer (Solarbio) and boiled at 95℃ for 5 min. Samples were shortly centrifugated before electrophoresis on polyacrylamide gels. Proteins were blotted on polyvinylidene fluoride (PVDF) membranes (0.22 μm, Millipore). Membranes were blocked by 5% BSA in TBST for 1 hr at room temperature and incubated with primary antibodies overnight at 4 ℃. After washing by TBST for 3 times, membranes were incubated with horseradish peroxidase (HRP) conjugated secondary antibodies (1:10,000, CST) for 1 hr at RT. Immunoblots were then developed using ECL Reagent (Millipore) and exposed by Amersham ImageQuant™ 800 (Cytiva). ImageJ was used to analyze the images.

### Lipid raft isolation

Cortical neurons were plated in 10 cm dish at 2 x 10^7^ /dish. After 5 days of culturing, the neurons were scraped off in 1 x TNE buffer (20 mM Tris-HCl(pH7.4), 250 mM sucrose, 1 mM MgCl_2_, 1mM CaCl_2_) with 1% Triton X-100 and protease inhibitor added before use. The cells were lysed by pipetting 15 times with a 1 mL pipette tip, followed by pipetting 10 times with a 26G needle. The lysates were centrifuged at 1000 x g for 10 min at 4℃ to remove the nuclei and incompletely lysed cells. 1.5ml of cell extracts were mixed with equal volume of Opti-Prep™, placed at the bottom of the ultracentrifuge tube (14 x 89 mm, Beckman), then overlaid with 2.5 ml of 30% Opti-Prep™, 2.5 ml of 20% Opti-Prep™ and 2 ml of 5% Opti-Prep™. Samples were then subjected to ultracentrifugation using SW41Ti rotor (Beckman) at 38,000 rpm (247,600 x g) for 18h at 4 ℃. Samples were divided into 12 fractions of equal volume collected from the top of the gradient and fraction 1-10 were then mixed with 2-fold volume of iced acetone and precipitate proteins for overnight at −20 ℃. The precipitations were resolved by 50μl 2% SDS buffer and subjected to western blot. For lipid raft isolation from hippocampal extracts, Kimble Dounce grinders were used to homogenize tissues. 3.75 mg proteins of each sample in 1.5 ml 1 x TNE buffer containing 1% Triton X-100 and protease inhibitor were subjected to ultracentrifugation using the same condition as cultured neurons but without acetone precipitation.

### Receptor internalization and immunohistochemistry

The internalization assay were performed based on a previous report [Yi, et al. Embo, 2020]. Briefly, cultured hippocampal neurons seeding on coverslips were first washed twice by ice-cold ACSF buffer (124 mM NaCl, 3.7 mM KCl, 1.0 mM MgSO4, 2.5 mM CaCl2, 1.2 mM KH2PO4, 24.6 mM NaHCO3, 10mM D-glucose). Neurons were then incubated with antibody against the extracellular domain of P75NTR (AF1157, R&D, 2μg/mL) diluted in ACSF buffer for 1 hr at 4 ℃ to label surface p75^NTR^. Following incubation, slips were washed in ACSF and incubated at 37 ℃ to allow internalization. At different time points, formic acid buffer was added to stop the internalization. Total staining (100%) was determined by fixing cells immediately after antibody feeding without acid wash step. The cells were fixed by 4% PFA for 15 min at room temperature and washed three times in PBS. For immunohistochemistry, cells were blocked in 5% NDS and 0.3% Triton X-100 for 15min at room temperature. Other primary antibody (RABs 1:400, Lamp1 1:100, MAP2 1:1000) except for p75^NTR^ were diluted in blocking buffer and incubated with cells overnight at 4 ℃. After incubation, cells were washed three times in PBS and proper secondary antibodies at 1:500 dilution were added to cell to incubate for 1h at RT. Cells were washed again in PBS and mounted for further visualization under confocal microscope.

### Immunohistochemistry

Cryosections (14 μm) of mouse brain were equilibrated at room temperature for 30min and washed twice in PBS buffer, Then sections underwent microwave-assisted antigen retrieval (500W, 5 min) in citrate buffer followed by gradual cooling. After cooling to RT, sections were washed for twice in PBS, and blocked in blocking buffer containing 5% NDS and 0.3% Triton X-100 for 30 min. Primary antibodies were diluted in blocking buffer, added to sections and incubated overnight at 4℃. Sections were washed three times by PBS and added with proper secondary antibodies for 1 hr at room temperature. 4′,6-Diamidino-2-phenylindole (DAPI, 0.5 μg/ml) was used to stain cell nucleus for 10min at RT after secondary antibody incubation. Sections were washed for 3 times by PBS. After washing, sections were mounted and examined with confocal microscope. For fixed neurons cultured on coverslips, same protocol was used but without equilibration and antigen retrieval.

### APP Fractionation and Aβ_1-42_ detection by ELISA

Cryosections (14 μm) of mouse brain were equilibrated at room temperature for 30min and washed twice in PBS buffer, Then sections underwent microwave-assisted antigen retrieval (500W, 5 min) in citrate buffer followed by gradual cooling. After cooling to RT, sections were washed for twice in PBS, and blocked in blocking buffer containing 5% NDS and 0.3% Triton X-100 for 30 min. Primary antibodies were diluted in blocking buffer, added to sections and incubated overnight at 4℃. Sections were washed three times by PBS and added with proper secondary antibodies for 1 hr at room temperature. 4′,6-Diamidino-2-phenylindole (DAPI, 0.5 μg/ml) was used to stain cell nucleus for 10min at RT after secondary antibody incubation. Sections were washed for 3 times by PBS. After washing, sections were mounted and examined with confocal microscope. For fixed neurons cultured on coverslips, same protocol was used but without equilibration and antigen retrieval.

### Barnes Maze

The Barnes maze spatial memory test was performed using 6-month old male mice. The maze platform was positioned 140 cm above the ground level, with a total diameter of 122 cm. Twenty evenly distributed openings (each 5 cm in diameter) were incorporated in the design. Spatial cues were placed around the maze, and a high-intensity light source was positioned directly above the maze as the stimulus. On the first day of training, the mouse was placed in the escape tunnel for 1 min(habituation). After habituation, the mouse was then placed at the center of the maze inside a dark box. The box was removed after 10s and the recording was on at the same time. The mouse was free to explore the maze for 3min or until the mouse escaped. If the mouse failed to find the tunnel, it would be put into the tunnel for 1 extra minute. The training was conducted once daily for 6 consecutive days. The platform was cleaned by 70% ethanol after individual exploration and was moved everyday by 90° to avoid any odorant cue. In the memory test sessions (short-term test: 2 hr after the last training; long-term test: 24 hr after the last training), the tunnel was removed and the mouse was allowed to explore the maze for 90 sec. Time spent in the target quadrant and exploration to the target opening was analyzed to evaluate the spatial memory of mouse.

### Statistical analysis

Statistical analysis were performed using GraphPad Prism 10 software. Results were presented as mean ± standard error of the mean (SEM). Student’s t-test, one-way ANOVA or two-way ANOVA were performed according to the requirements of the experiment. Statistical significance: *p<0.05; **, p<0.01; ***, p<0.001.

## Acknowledgements

The authors would like to thank Lei Wang, Jocelyn Jia, Yankui Fu and Shuo Zhang for technical and admin assistance. This work was supported by research grants to C.F.I. from Peking University, Chinese Institute for Brain Research, Beijing, and Swedish Research Council (Vetenskapsrådet, contract nr. 2024-03222); and a startup grant to M.X. from Swedish Research Council (Vetenskapsrådet, contract nr. 2021-01805).

## Author contributions

Y.M. conceived the project, performed all experimental work, analyzed data and prepared a draft of the manuscript and figures; M.X. co-directed the project and corrected the manuscript; C.F.I. directed the research and wrote the final version of the manuscript.

## Supplementary figure legends

**Supplementary Figure S1.**
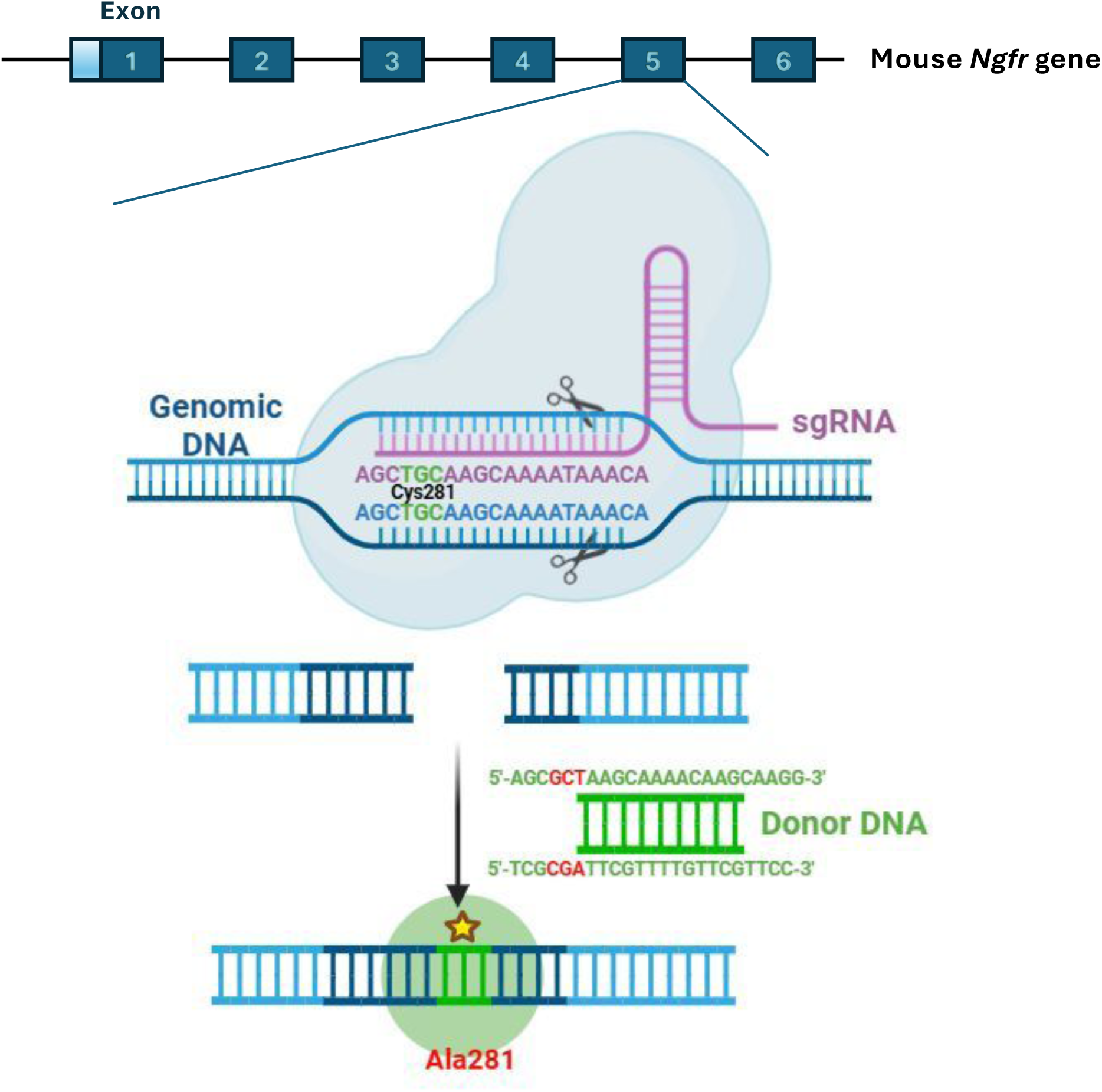
Schematic of strategy for generating palmitoylation-deficient p75^C281A^ mutant mice.

**Supplementary Figure S2.**
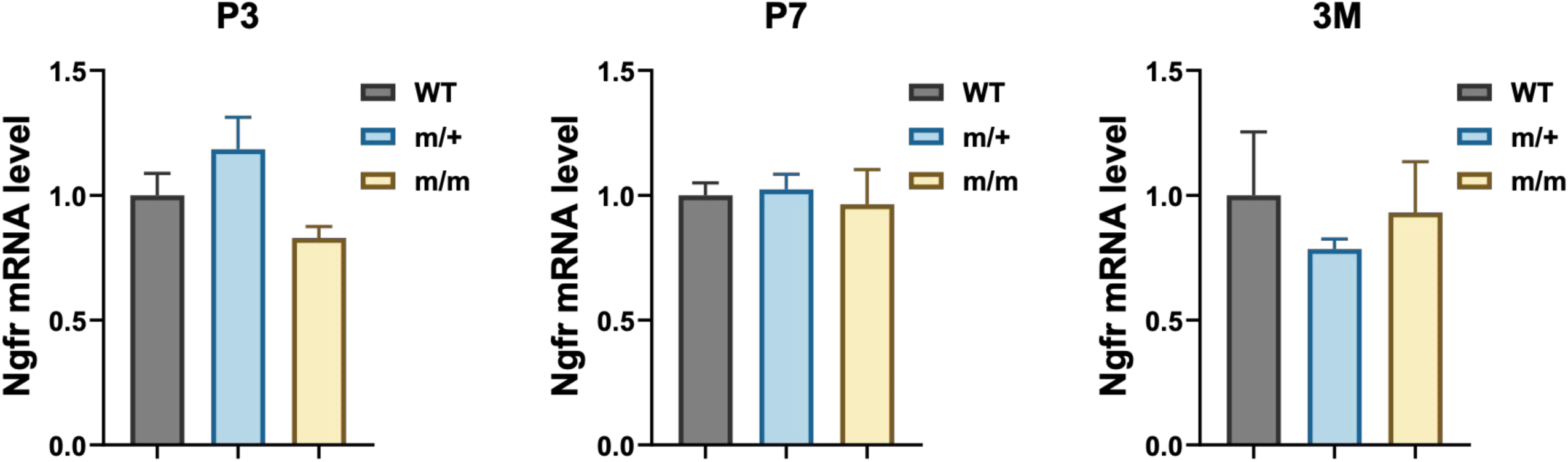
p75^NTR^ mRNA levels are not affected by the Cys^281^Ala mutation. Quantitative PCR analysis of p75^NTR^ (*Ngfr*) mRNA levels in the hippocampus of wild type (WT), mutant heterozygous (m/+) and homozygous (m/m) p75^NTR^ mice at postnatal day 3 (P3), P7, and 3 months (3M) of age. Statistical analysis by one-way ANOVA; mean ± SEM; N=3 animals per group at each stage.

**Supplementary Figure S3.**
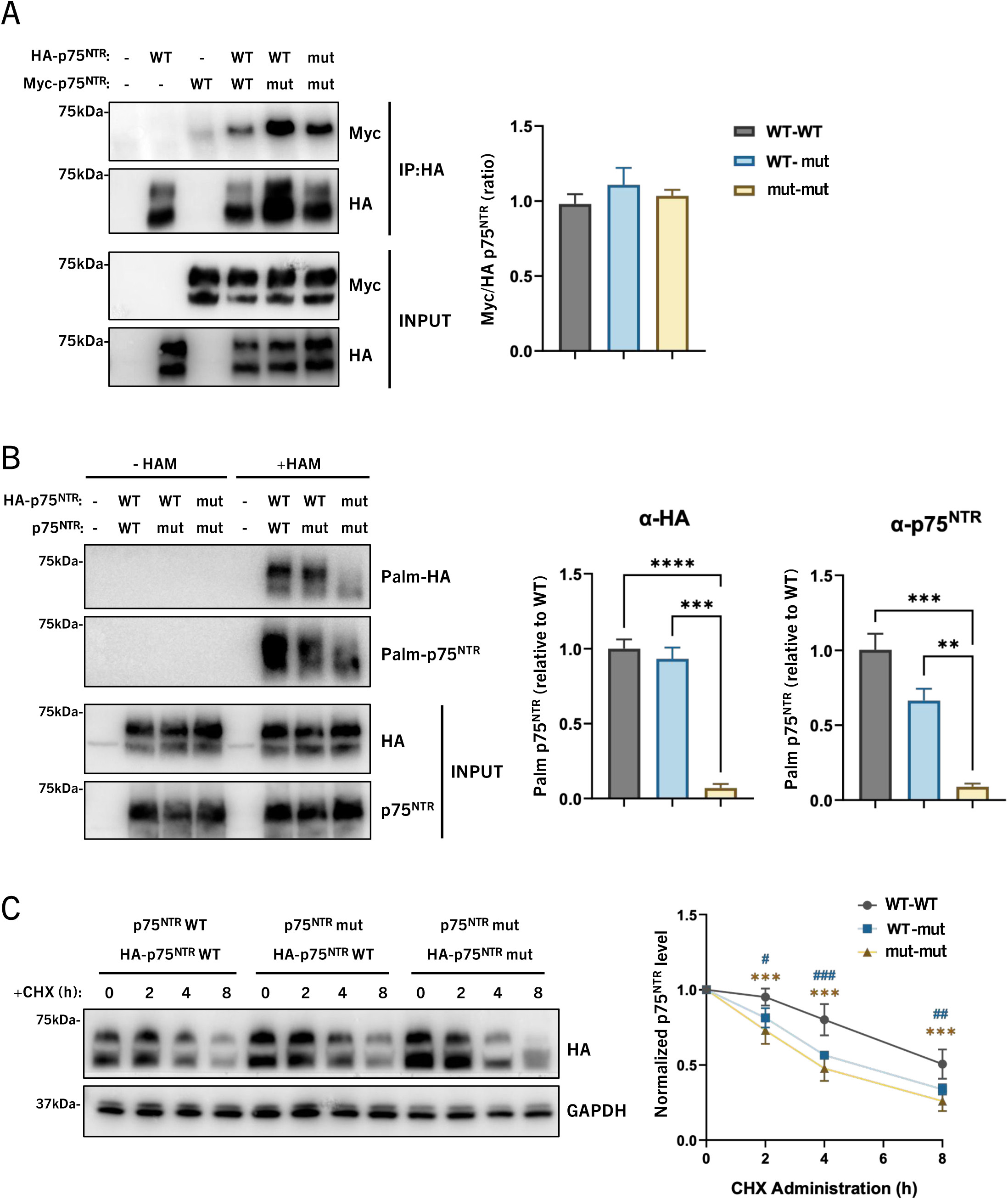
Dominant effect of palmitoylation-deficient p75^NTR^ on the half-life of wild type p75^NTR^. (A) Lack of palmitoylation does not interefere with p75^NTR^ dimerization. Co-immunoprecipitation between HA-tagged p75^NTR^ (wild type [WT] or palmitoylation-deficient p75^NTR^ mutant [mut]) with Myc-tagged p75^NTR^ (WT or mut) in HEK293T cells that co-transfected with expressing plasmids for 24h. Statistical analysis by one-way ANOVA; mean ± SEM; N=4 independent experiments. (B) Lack of palmitoylation does not interefere with the palmitoylation of the interacting wild type protomer in the p75^NTR^ dimer. ABE assay analysis of the palmitoylation level of wild type (WT) or palmitoylation-deficient p75^NTR^ mutant (mut) HA-tagged p75^NTR^ when co-expressed with homogenous or heterogenous p75^NTR^ without tag. Histogram shows the relative palmitoylation level of HA-tagged p75^NTR^ (left panel) and all p75^NTR^ (right panel) compared to the condition in which only wild type p75^NTR^ is expressed. Statistical analysis by one-way ANOVA; mean ± SEM; N=3 independent experiments; **p<0.01, ***p<0.001, ****p<0.0001. (C) Reduced half-life of wild type p75^NTR^ upon co-expression of palmitoylation-deficient mutant. Cycloheximide (CHX) pulse-chase assay of half-life of HA-tagged p75^NTR^ when co-expressed with wild type (WT) or palmitoylation-deficient mutant p75^NTR^ without tag. Statistical analysis by two-way ANOVA followed by Tukey’s multiple comparisons test; mean ± SEM; N=5 independent experiments; ***p<0.001 m/m(m/m) versus WT(WT); #p<0.05, ##p<0.01, ###p<0.001 WT(m/m) versus WT(WT).

**Supplementary Figure S4.**
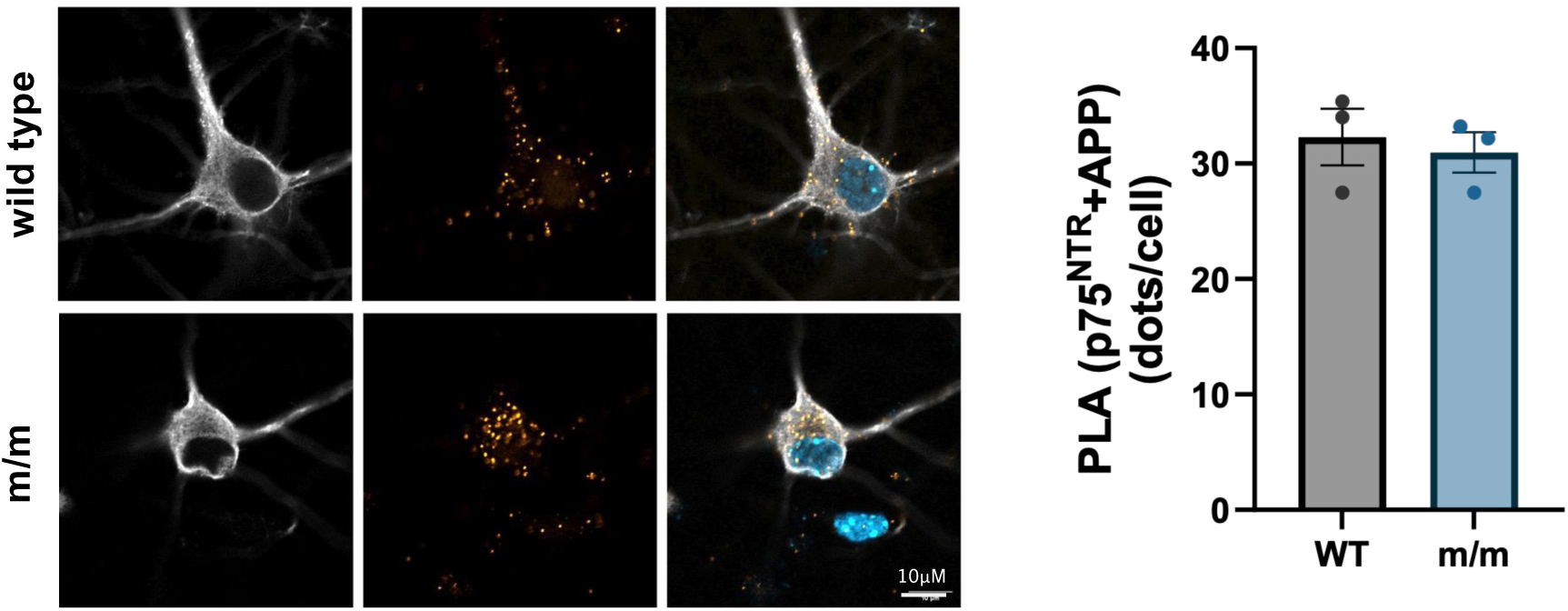
Unchanged APP localization to lipid rafts in hippocampus of 5xFAD mice expressing palmitoylation-deficient p75^NTR^. (A) Western blot analysis of p75^NTR^ and lipid raft markers caveolin-1 (CAV-1) and ganglioside GM1 in lipid raft fractions 1 to 10 from cultured cortical neurons. Fractions 11 and 12 were excluded due to saturating levels of p75^NTR^ protein. Extremely low levels (<0.1%) of p75^NTR^ protein can be detected in fractions 3 and 4 corresponding to lipid rafts. (B) Western blot analysis of APP and lipid raft marker caveolin-1 (CAV-1) in lipid raft fractions 1 to 12 from total extracts isolated from the hippocampus of 9 month old 5xFAD mice. Very low levels (<1%) of p75^NTR^ protein can be detected in fractions 5 and 6 corresponding to lipid rafts. (C) Western blot analysis of APP in lipid raft fractions 5 and 6 (marked by CAV-1) isolated from total extracts of 9 month old 5xFAD mice expressing wild type p75^NTR^ (5xFAD) or palmitoylation-deficient p75^NTR^ (m/m;FAD). Unfractionated total extracts (input) are also shown for comparison. Results from two independent experiments (batch 1 and 2) are shown. (D) Quantitative analysis of APP levels in lipid raft fractions (as shown in panel C) normalized to levels in 5xFAD (expressing wild type p75^NTR^).

**Supplementary Figure S5.**
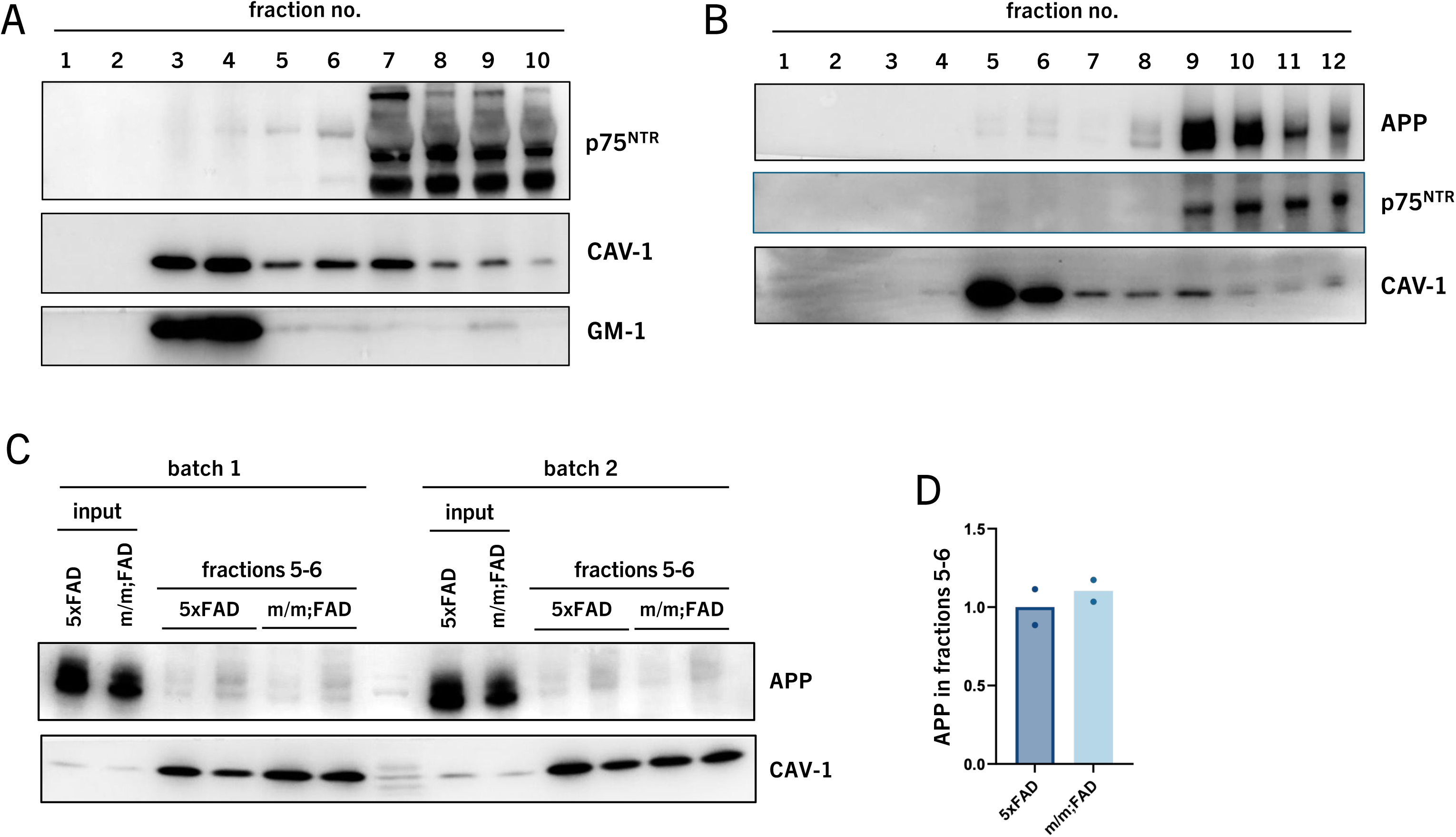

